# Structural Insights into Bifunctional Thaumarchaeal Crotonyl-CoA Hydratase and 3-Hydroxypropionyl-CoA Dehydratase from *Nitrosopumilus maritimus*

**DOI:** 10.1101/2021.09.22.461329

**Authors:** Ebru Destan, Busra Yuksel, Bradley B. Tolar, Esra Ayan, Sam Deutsch, Yasuo Yoshikuni, Soichi Wakatsuki, Christopher A. Francis, Hasan DeMirci

## Abstract

The ammonia-oxidizing thaumarchaeal 3-hydroxypropionate/4-hydroxybutyrate (3HP/4HB) cycle is one of the most energy-efficient CO_2_ fixation cycles discovered thus far. The protein encoded by Nmar_1308 (from *Nitrosopumilus maritimus* SCM1) is a promiscuous enzyme that catalyzes two essential reactions within the thaumarchaeal 3HP/4HB cycle, functioning as both a crotonyl-CoA hydratase (CCAH) and 3- hydroxypropionyl-CoA dehydratase (3HPD). In performing both hydratase and dehydratase activities, Nmar_1308 reduces the total number of enzymes necessary for CO_2_ fixation in Thaumarchaeota, reducing the overall cost for biosynthesis. Here, we present the first high-resolution crystal structure of this bifunctional enzyme with key catalytic residues in the thaumarchaeal 3HP/4HB pathway.

## Introduction

CO_2_ fixation allows autotrophic organisms to use inorganic carbon and energy acquired from light or chemical reactions as a means to synthesize the entirety of their biomass^1^. This process is dually important: in addition to biosynthesis, it also serves a role in the removal of CO_2_ from the environment. Humans have significantly altered the CO_2_ concentrations present in the Earth’s atmosphere through industrialization and burning of fossil fuels^2^. In order to curb this increase in atmospheric CO_2_ levels, improvements must be made in sequestration methods and energy efficiency^3^. Studying the means by which microorganisms have adapted their carbon fixation pathways and metabolisms to be more efficient in low-energy/nutrient environments offers a possible solution to this problem.

Archaea belonging to the phylum Thaumarchaeota (formerly known as Marine Group I Crenarchaeota) are well adapted to life in low nutrient environments^1,4,5^ such as the open ocean, where they are estimated to comprise over 30 % of planktonic bacterial and archaeal populations^6^. Over 15 years ago, Thaumarchaeota were first discovered and shown to play a key role in the global nitrogen cycle - oxidizing ammonia to nitrite as the first step in nitrification - through molecular/genomic surveys and the subsequent isolation of *Nitrosopumilus maritimus*^7,8,9,10^. Since that time, Thaumarchaeota have been found in nearly every environment on Earth, and often have higher abundances than their ammonia-oxidizing bacteria (AOB) counterparts^11,12,13,14^. Marine Thaumarchaeota are believed to thrive in nutrient-limited environments due to specific adaptations^4,5^, requiring only nano-molar level ammonia^15,16^. This provides an advantage for ammonia-oxidizing archaea (AOA), as they are in competition for the same limited resources as AOB and other microorganisms in shared environments^17,18^.

All known ammonia-oxidizing Thaumarchaeota fix CO_2_ with a modified version of the 3HP/4HB cycle originally described in Crenarchaeota and further characterized in *N. maritimus*^1,19,20^**(Fig. 1)**. This cycle can be broken down into two characteristic halves: (1) In the first part of the cycle, one acetyl-CoA and two bicarbonate molecules are converted to succinyl-CoA through 3-hydroxypropionate intermediate; (2) this is followed by the conversion of succinyl-CoA to regenerate acetyl-CoA, producing two molecules of acetyl-CoA by way of a 4-hydroxybutyrate intermediate. From here, one of the acetyl-CoA molecules generated then serves as a carbon precursor for the next round of the cycle^1,21^. The individual steps of this cycle, however, distinguish it from other known aerobic and autotrophic CO_2_ fixation pathways, as well as provide further evidence of the differentiation and efficiency of the 3HP/4HB cycle used by Thaumarchaea versus their Crenarchaeal ancestors^1^. Notably, the thaumarchaeal 3HP/4HB cycle requires two fewer high-energy phosphoric anhydride bonds of ATP per acetyl-CoA molecule than the crenarchaeal 3HP/4HB cycle and three fewer than the Calvin-Benson-Bassham cycle^1^.

**Figure 1.**
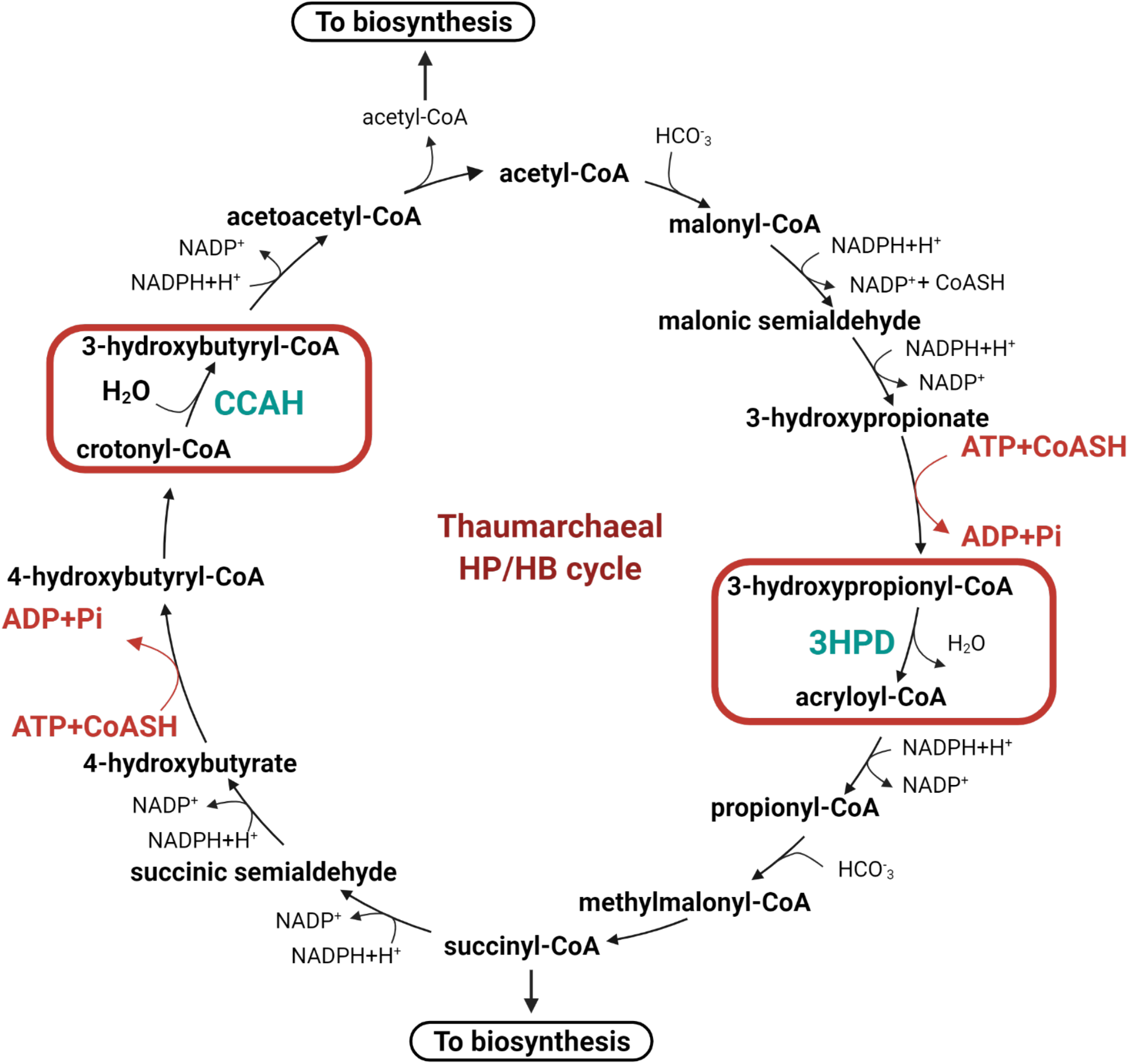
The dual enzymatic steps catalyzed by Nmar_1308 in the thaumarcheal 3HP/4HB cycle. Representation of 3HP/4HB cycle reactions in the thaumarcheal variant. The reaction showing the energy efficiency of the taumarcheal cycle is colored in red. Highlighted rectangle in red indicates the bifunctional reactions catalyzed by the Nmar_1308 protein. Nmar_1308 is a promiscuous enzyme with dual functions that dehydrates 3-hydroxypropionyl-CoA to acryloyl-CoA via 3-hydroxypropionyl-CoA dehydratase (3HPD) and hydrates crotonyl-CoA to 3-hydroxybutyryl-CoA via crotonyl-CoA hydratase (CCAH). 3HPD and CCAH are colored teal.

By breaking down the 3HP/4HB cycle into individual steps, we can focus on the enzymes responsible for catalyzing each respective reaction. In doing so, the structure and kinetics of each enzyme can be determined to provide more insight into how the overall 3HP/4HB cycle in Thaumarchaeota is so energy efficient. Here we have focused on the Nmar_1308 enzyme because it plays an essential bifunctional role in the thaumarchaeal 3HP/4HB cycle, functioning as both as a 3-hydroxypropionyl-CoA dehydratase (3HPD) and crotonyl-CoA hydratase (CCAH)^1,22^. In its capacity as a 3HPD, Nmar_1308 catalyzes the conversion of 3-hydroxypropionyl-CoA to acryloyl-CoA and water. Conversely, as a CCAH Nmar_1308 converts crotonyl-CoA to 3-hydroxybutyryl-CoA through a dehydration reaction. The promiscuity of Nmar_1308 and a second bifunctional enzyme in the *N. maritimus* 3HP/4HB cycle (acetyl-CoA / propionyl-CoA carboxylase) reduces the cost of protein biosynthesis as another means to increase overall efficiency^1^. In addition, it was recently discovered that the Nmar_1308 homolog in the crenarchaeon *Metallosphaera sedula* (5ZA1; Msed_2001), previously characterized as 3HPD^23^, can also function as a CCAH^22,23^.

Despite the critical role of Nmar_1308 protein in this aerobic archaeal CO_2_ fixation pathway, high-resolution structural details and dynamics of this novel enzyme remain unknown. Here, we determined the first high-resolution crystal structure of Nmar_1308, in order to better understand the role this enzyme plays in the thaumarchaeal modification of the autotrophic 3HP/4HB cycle that allowed it to become one of the most energy-efficient CO_2_ fixation pathways known.

## Results & Discussion

### Apo-form Nmar_1308 structure

The Nmar_1308 gene encodes a bifunctional protein that plays critical roles in the 3HP/4HB cycle by functioning both as 3HPD and CCAH^1,22^ (**Fig. 1**). The apo-form structure of Nmar_1308 was determined as hexameric form at 2.15 Å resolution in space group *P*3_2_ **(Fig. 2 & Table 1)**. The homohexameric structure consists of a dimer of trimers arrangement. The asymmetric unit contains 6 Nmar_1308 monomers with a total of 1422 residues. Chain A has 232 modeled residues while chain B has 236; chain C has 237; chain D&E have 230 and chain F has 229 modeled residues. Each subunit contains an active site with catalytic residues Gly109, Glu112 and Glu132 and none of the chains have bound substrates/products or metal ions. These three residues are a part of the well documented canonical active site of bacterial and archaeal enoyl-CoA hydratase/dehydratases, involving two glutamate residues and two backbone amides forming the oxyanion hole (**Fig. 3A-C**).

**Table 1.** Data collection and refinement statistics of X-ray crystallography data collection and structure refinement.

|  |  |
| --- | --- |
| Dataset | NMAR_1308 |
| PDB ID | 7EUM |
| **Data collection** |  |
| Beamline | SSRL BL-12-2 |
| Space group | P 32 |
| Cell dimensions<br>*a, b, c* (Å) | 75.87, 75.87, 207.96 |
| $\alpha, \beta, \gamma$ (°) | 90, 90, 120 |
| Resolution (Å) | 37.93-2.15 (2.28-2.15) |
| *R*<sub>merge</sub> (%) | 12.7 (188.3) |
| *CC1/2* | 99.9 (52.2) |
| *I* / $\sigma I$ | 13.38 (1.22) |
| Completeness (%) | 99.7 (98.8) |
| Redundancy | 10.51 (10.49) |
| **Refinement** |  |
| Resolution (Å) | 29.69-2.15 (2.21-2.15) |
| No. reflections | 72674 (5907) |
| *R*<sub>work</sub> / *R*<sub>free</sub> | 0.24/0.28 (0.30/0.31) |
| No. atoms |  |
| Protein | 10440 |
| Ligand/Ion/Water | 336 |
| *B*-factors |  |
| Protein | 64.6 |
| Ligand/Ion/Water | 51.3 |
| Coordinate errors | 0.32 |
| R.m.s deviations |  |
| Bond lengths (Å) | 0.002 |
| Bond angles (°) | 0.562 |
| Ramachandran plot |  |
| Favored (%) | 96.08 |
| Allowed (%) | 3.85 |
| Disallowed (%) | 0.07 |

Numbers corresponding to the highest resolution shell are shown in parentheses.

**Figure 2.**
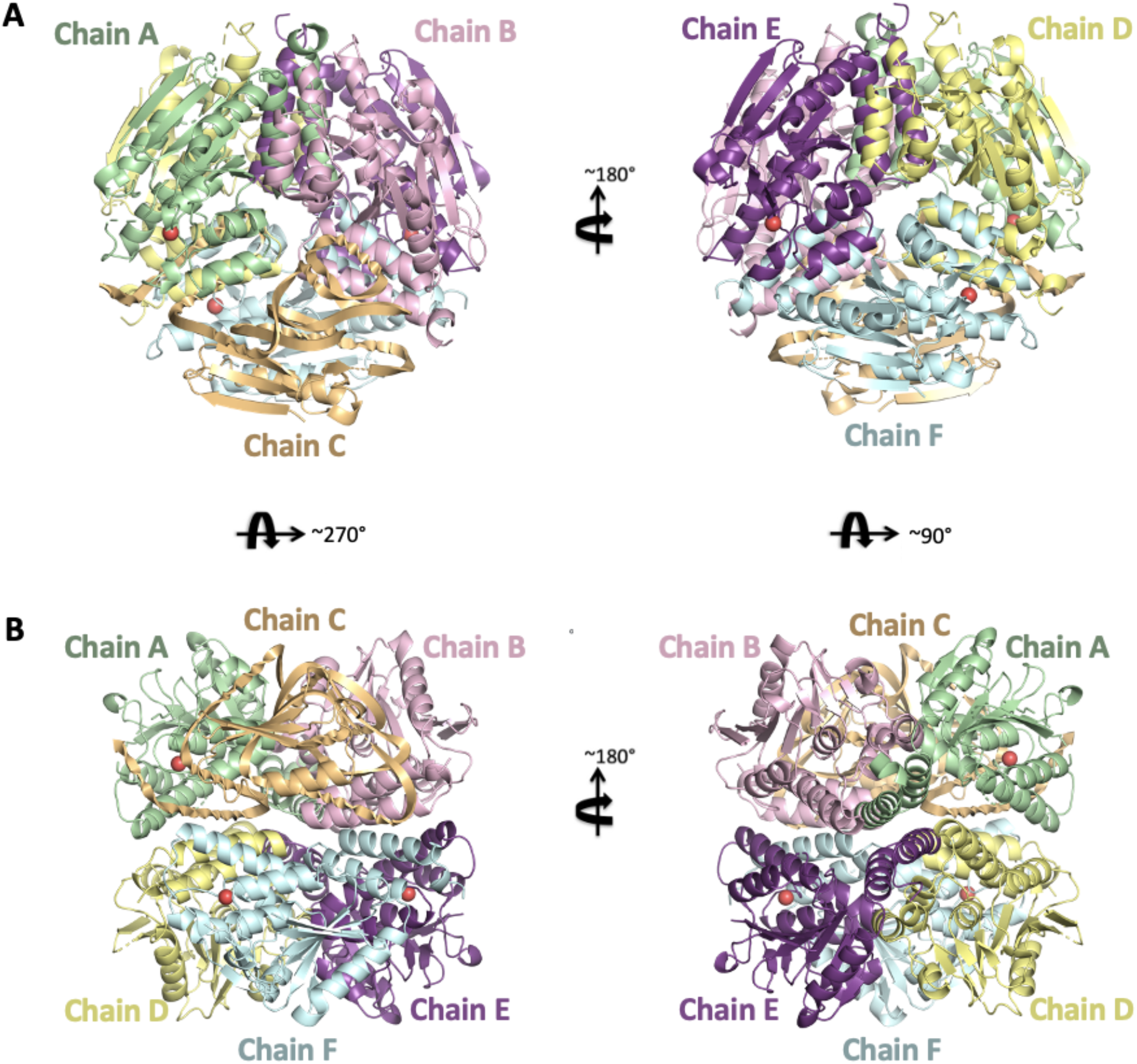
Cartoon representation of Nmar_1308 structure. **(A)** The overall structure of Nmar_1308 with the upper view is colored based on each chain. **(B)** The overall structure of Nmar_1308 with the side view is colored based on each chain. Chains are colored in pale green, light pink, light orange, pale yellow, violet purple and pale cyan, respectively. The critical water molecules which are present in chains A, E and F are indicated as red spheres.

**Figure 3.**
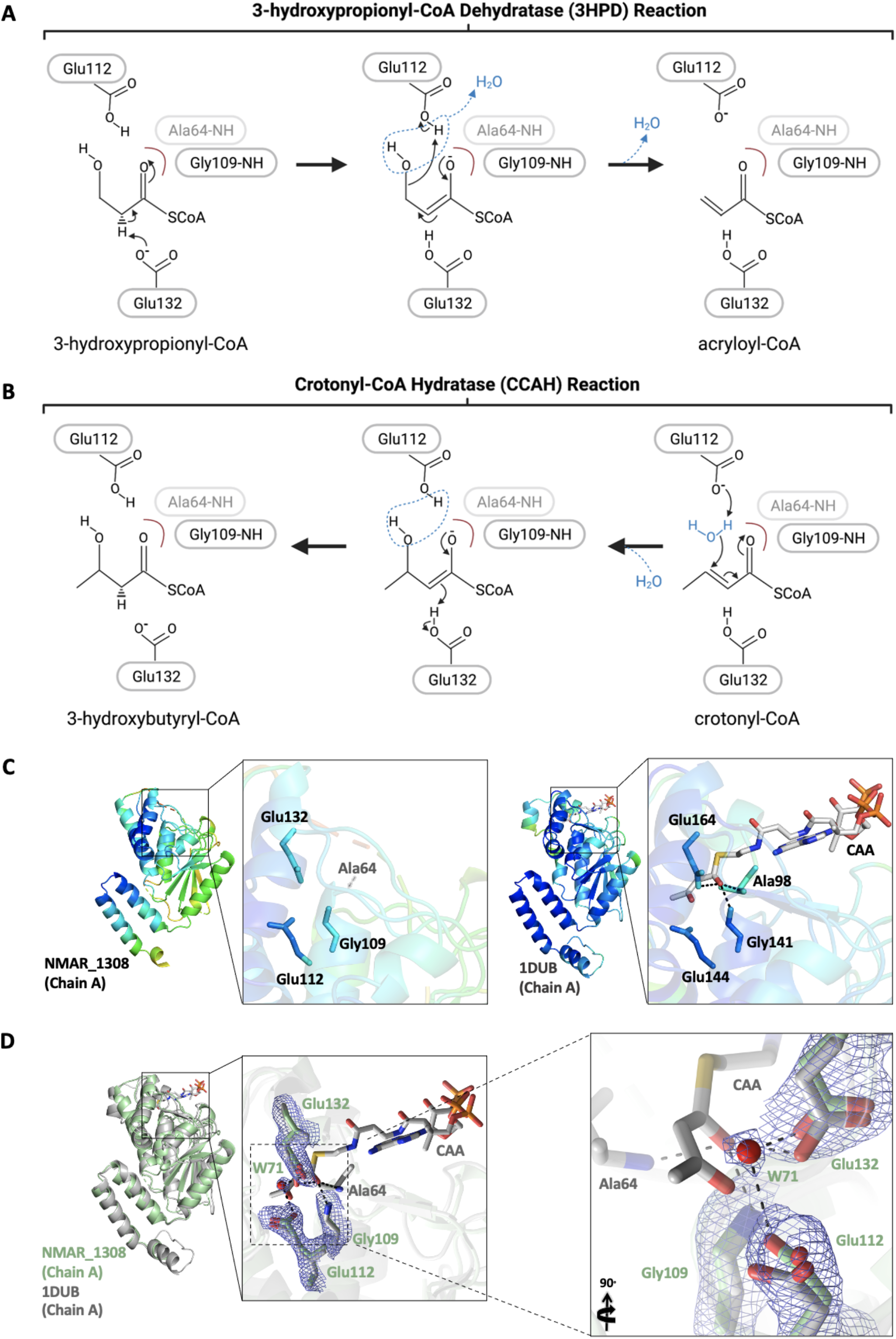
Representation of hydratase/dehydratase active sites. The proposed mechanism of hydratase **(A)** and dehydratase **(B)** reactions by Nmar_1308. One of the two residues forming the oxyanion hole for the catalyst, Ala64 is missing in our structure, but is indicated in faint gray at the putative position. **(C)** The ligand binding pocket is shown for Nmar_1308 structure and the 2-enoyl-CoA hydratase structure (PDB ID: 1DUB) to compare the stability around the binding pocket. The ligand of 1DUB structure, acetoacetyl-CoA (CAA) is colored in gray. **(D)** Superposition of the apo-form Nmar_1308 protein and the structure of rat liver mitochondrial enoyl-CoA hydratase complexed with acetoacetyl-CoA. The Nmar_1308 structure is colored in pale green while the superposed enoyl-CoA hydratase structure (PDB ID: 1DUB) is colored in gray. Hydrogen bond interactions are shown with dashed lines in black. The 2Fo-Fc electron density of the Nmar_1308 is contoured at 1 σ level and colored in slate.

### Water molecule between catalytic residues at the active site

Two catalytic residues Glu112 and Glu132 play key roles during catalysis, one as an acid and the other as a base. In the Nmar_1308 structure, they form hydrogen bonds with a water molecule (W71) (**Fig. 3D**). Two other chains in the hexamer have water molecules at the same position, chain E (W188) and chain F (W239), while the other three are empty. The position of these water molecules (W71, W188, and W239) are very close to the water positions bridged by the corresponding two glutamic acid side chains of the other hydratase structures^24,25^. This addition of the water to the C3 catalyzes the hydration reaction from crotonyl-CoA to 3-hydroxybutyryl-CoA, and the dehydratase reaction performs in the reverse direction from 3-hydroxypropionyl-CoA to acryloyl-CoA by removal of water (**Fig. 3A**). In the hydratase reaction, Glu132 protonates C2 of a ligand while Glu112 activates the water molecule that is to be added to C3 from the other side of the substrate^24,26^. In the reverse reaction, Glu112 acts as an acid attacking the hydroxyl oxygen of C3 for removal of water assisted by Glu132 as a base to extract a proton from C2. The hexamer of Nmar_1308 with three monomers having the water and the other three without may indicate the flexibility of the enzyme in accommodating either substrate for hydratase or dehydratase reactions.

### Dynamic oxyanion hole

The apo-form Nmar_1308 crystal structure lacks electron density for residues 62-69 (some monomers lack one or two fewer residues) in 6 monomers, indicating the flexible nature of this loop located on the outer surface of each monomer. This loop is visible in ligand-bound structures of the related hydratases and dehydratases, indicating that the loop is flexible without a bound ligand. The residues near this loop display high B factors in both the apo-form Nmar_1308 and the inhibitor-bound structure of enoyl-CoA hydratase from *Rattus norvegicus* (PDB ID: 1DUB) **(Fig. 3C)**. In contrast, the core of the enzyme hexamer is rather rigid including the regions in and around the binding pocket for the enoyl-group of substrates **(Fig. 3C)**. In the proposed hydratase/dehydratase reaction mechanisms, two conserved residues alanine and glycine (Ala64 and Gly109 in Nmar_1308) in the binding pocket act as an oxyanion hole by offering their backbone amides to the carbonyl oxygen (O_1_) of the substrate and stabilizing the negative charge accumulated on O1 oxygen in the transition state. Interestingly, Ala64 is part of the abovementioned flexible loop disordered in the apo-form Nmar_1308 structure while Gly109 is at the N-term end of a relatively rigid α3 helix. Again, in the holo-form of the hydratase and dehydratase structures, the conserved alanine of the oxyanion hole is stable. Taken together, the oxyanion hole is dynamically assembled when a substrate is bound at the active site and disassembled as the product leaves.

### Comparison with other hydratase/dehydratase structures and substrate specificity

To further confirm these notions of the active site architecture and flexibility, we then expanded the structure comparison to representative homologous structures **(Fig. 4)**. Superposition of the Nmar_1308 structure with 11 available structures indicates that there are minor conformational differences between Nmar_1308 protein and homologous hydratase/dehydratases structures. These structures were classified based on their catalytic properties (hydratase vs. dehydratase) or their complex states (apo or holo). Among these structures, hydratases in holo form (PDB IDs: 1MJ3, 1DUB, 2DUB, 1EY3, 3Q0G) and in apo form (PDB IDs: 3PZK, 3Q0J, 3H81, 5XZD); and a dehydratase in complex with its ligand (holo form) were (PDB IDs: 5JBX) superposed with the Nmar_1308 structure with the indicated root mean squared deviation (RMSD) values. Msed_2001 (PDB IDs: 5ZAI) is shown as bifunctional^22,26^. It is notable that the RMSD value between Msed_2001 and Nmar_1308 is quite small, 0.91 Å, the lowest among the hydratase/dehydratase compared in this study (the other RMSD are 1.04 Å to 1.22 Å). To provide a better understanding of these conformational differences and highlight the importance of these specific residues (which may define Nmar_1308 as a bifunctional enzyme), a multiple sequence alignment was performed using Jalview^27^ **(Fig. 5)**. The structure-based sequence alignment analysis shows the residues critical for the hydratase/dehydratase reactions, Ala64, Gly109, Glu112 and Glu132 are well conserved **(Fig. 5A&B)**, and consistent with the active site residues reported earlier^26,28,29^.

**Figure 4.**
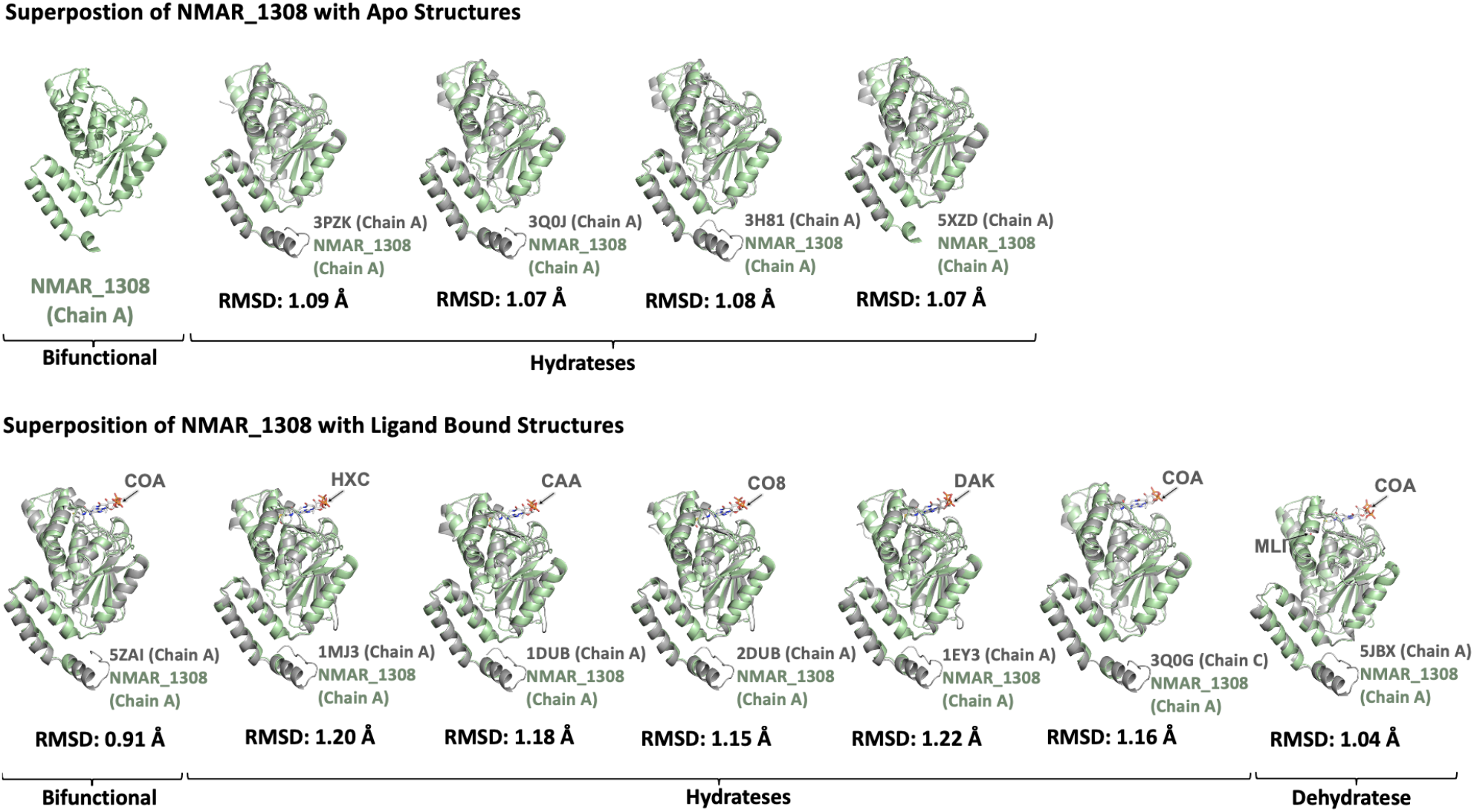
Superposition of Nmar_1308 with homologous structures. Superposition is performed by using cryo-synchrotron structure of Nmar_1308 with published apo/ligand bound structures of enoyl-CoA hydrates and 3-hydroxypropionyl-CoA dehydratase (3HPD). Average RMSD values were calculated from all pairwise RMSD values between Cα atoms of an Nmar_1308 monomer and one of the superimposed monomers of homologous enzymes (36 pairs for hexamers and 18 for trimers), and are indicated below each panel. All standard errors of the RMSDs are below 0.01 Å. Chain A of Nmar_1308 is colored in pale green while the superposed structure is colored in gray. The ligands were represented with arrows. (**HXC:** hexanoyl-CoA, **CAA:** acetoacetyl-CoA, **CO8:** octanoyl-CoA, **DAK:** 4-(N,N-dimethylamino)cinnamoyl-CoA, **COA:** coenzyme A)

**Figure 5.**
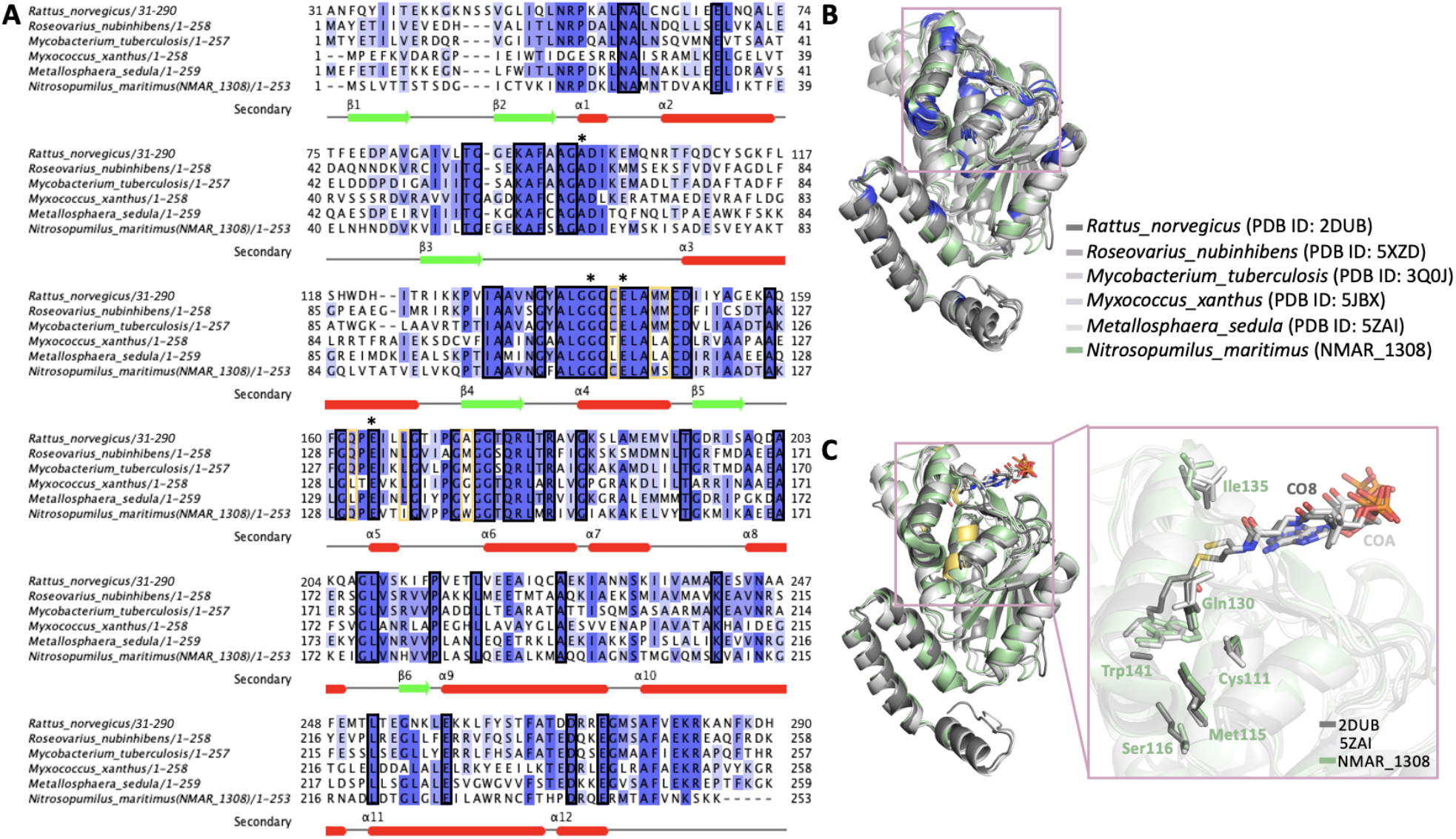
Overall representation of sequence alignment results. **(A)** Homologous hydratases (*Rattus norvegicus* (2DUB), *Roseovarius nubinhibens* (5XZD), *Mycobacterium tuberculosis* H37Rv (3Q0J)) and dehydratase (*Myxococcus xanthus* (5JBX)), bifunctional (*Metallosphaera sedula* (5ZAI)) sequences are aligned to show conserved and critical residues for functioning of Nmar_1308. Black squares represent the conserved residues in the sequence alignment while yellow orange color is used to indicate the observed critical residues. The catalytic residues during hydratase/dehydratase reaction is indicated with asterisks on the top of the sequence. **(B)** Slate color indicates conserved regions on the structure based on the sequence alignment result in panel A. The detected conserved residues in the hydrophobic binding pocket are shown with a pink square. **(C)** Residues implicated in the catalysis, Cys111, Met115, Ser116, Gln130, Ile135, and a residue important for limiting substrate size, Trp141, around the hydrophobic binding pocket are shown on the superposed structures of Nmar_1308, *Rattus norvegicus* hydratase (PDB ID: 2DUB) and bifunctional dehydrate/hydratase from *Metallosphaera sedula* (PDB ID: 5ZAI).

The structure-based multiple sequence alignment shows how Nmar_1308 restricts the range of short enoyl-CoA and 3-hydroxy-acyl-CoA substrates using Trp141 in the enoyl-CoA binding pocket. In other short enoyl-CoA hydratases this residue is Tyr142 in 3HPD of *M. sedula* (PDB ID: 5ZAI)^26^. These bulky residues serve as a limiter of the substrate length by closing the bottom of the substrate enoyl-CoA binding pocket. In contrast, smaller or thinner side chains such as alanine (*Rattus norvegicus* (2DUB)), methionine (*Roseovarius nubinhibens* (5XZD)), *Myxococcus xanthus* (5JBX)), and glycine (*Mycobacterium tuberculosis* H37Rv (3Q0J)) can allow for longer chain enoyl-CoA as substrates. In addition, Trp231 provides additional hydrophobic constriction from helix α11 of the neighboring monomer, hence the trimer is the minimal biological enzyme unit for hydratase and dehydratase functions. This is well conserved between long and short chain enoyl-CoA hydratases/dehydratases; tryptophan, phenylalanine, or tyrosine. These residues play important roles to define substrate specificity.

### Extended hydrogen bond network at the active site of dual function hydratase/dehydratase

The multiple sequence alignment also suggests that there are several less well conserved residues: Cys111, Ser116, and Gln130 **(Fig. 5A&C)**. They might play a role for assisting the bifunctionality of this protein, as they are in the vicinity of the active site of the hydratase/dehydratase **(Fig. 6 & Table 2)**. For example, while Cys111 is conserved in homologous hydratase structures^29,30^, dehydratase structures show either threonine or leucine^26,31^. The sulfur atom of Cys111 is actually very close to side chain oxygen of Gln130 (distance of 3.5 A). In the structures where Cys111 is replaced with leucine, the residue corresponding to Gln130 is replaced by another leucine, hence forming a hydrophobic interaction. These Cys111-Gln130 or Leu-Leu/Thr pairs are probably the results of co-evolution of these two residues to keep the stable active site architecture **(Fig. 6 & Table 2)**. Another residue which shows variation is Ser116 that is one turn upstream of the catalytically important Glu112 along α3 helix, partnering in the hydrogen bond network with Glu112. The serine is replaced by a methionine in homologous hydratase structures and alanine in the homologous dehydratase structure, both of which may not be able to participate in the hydrogen bond network of Glu112. Gln130 is near the other glutamic acid Glu132 with a hydrogen bond between their oxygen atoms, hence participating indirectly in the hydratase/dehydratase reaction. This glutamine is conserved in homologous hydratase structures while both dehydratase and bifunctional enzyme have a leucine (Leu130 for dehydratase from *Myxococcus xanthus* (5JBX), and Leu 131 for the bifunctional enzyme from *Metallosphaera sedula* (5ZAI)). These leucine residues will not be able to participate in the hydrogen bond network.

**Table 2.** The residues implicated in the dual-function based on multiple sequence alignment in Figure 5A.

|  |  |  |  |  |
| --- | --- | --- | --- | --- |
| Dual function<br>(NMAR_1308) | Cys111 | Ser116 | Gln130 | Panel A |
| Hydratase<br>(5JBX) | Cys | Met | Gln | Panel B |
| Dehydratase<br>(5JBX) | Thr | Ala | Leu | Panel C |
| Dual function<br>(5ZAI) | Leu | Ala | Leu | Panel D |

**Figure 6.**
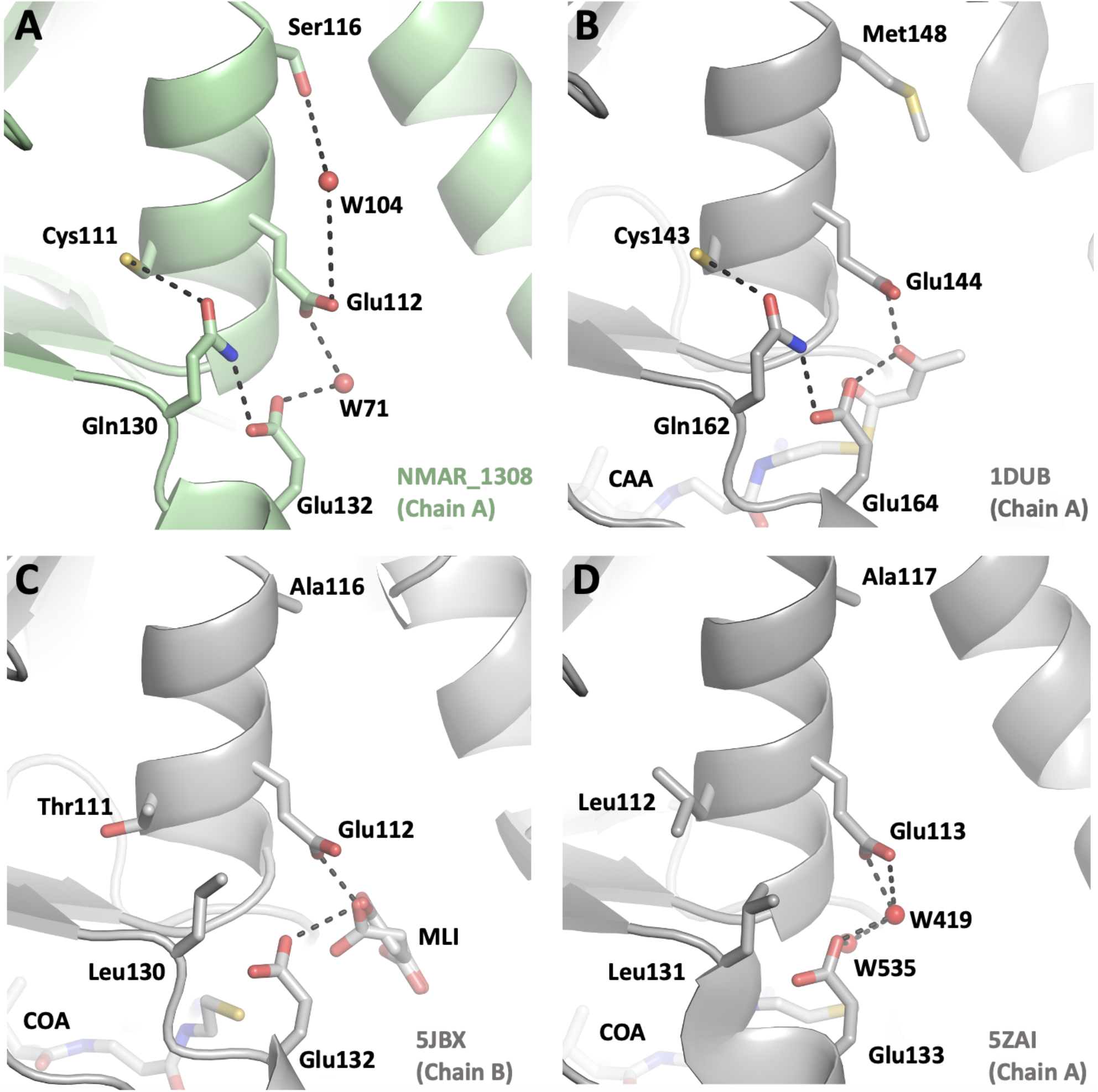
Representation of the potential interaction network of Nmar_1308 based hydratase/dehydrate structures. **(A)** Chain A of Nmar_1308 is indicated with a close look to detect the interaction partners during hydratase/dehydrates reaction. Chain A of Nmar_1308 is colored in pale green. **(B)** The hydratase structure from *Rattus norvegicus* (PDB ID: 1DUB) is colored in light gray. **(C)** The dehydratase structure from *Myxococcus xanthus* (PDB ID: 5JBX) is colored in light gray. Malonate (MLI) which may act as a water molecule by forming hydrogen bonds with two catalytic residues (Glu132 and Glu112) in the ligand bound structure is presented with sticks in the panel C. **(D)** The bifunctional enzyme from *Metallosphaera sedula* (PDB ID: 5ZAI) which has both hydratase and dehydratase activity is colored in light gray.

In summary, this study has provided the first high-resolution crystal structure of an enzyme within the thaumarchaeal 3HP/4HB cycle. Thaumarchaeota not only represent one of the most abundant microbial groups on the entire planet, but this CO_2_ fixation pathway is one of the most energy-efficient pathways known. Thus, the molecular insights into the bifunctional Nmar_1308 enzyme obtained in this study have implications for understanding the global C cycle (e.g., dark CO_2_ fixation) as well as for bioengineering applications using synthetic biology approaches.

## Methods

### Cloning

Initially, the Nmar_1308 gene was cloned into various expression vectors for expression tests by the JGI Synthetic Biology Group. After finding the optimal expression vector, the Nmar 1308 gene construct was purchased from Genscript Biotech (codon-optimized with cleavable N-terminal hexa-Histidine tag). Within the gene, NdeI and BamHI endonuclease restriction sites were used to insert Nmar_1308 into the pET28a vector plasmid. The plasmid was transformed into E. *coli*, strain BL21(Rosetta-2), from EMD Millipore. Transformed *E. coli* were then grown overnight on agar plates (50 µg/ml kanamycin, 30 µg/ml chloramphenicol) at 37 °C.

### Protein Expression and Purification

Overnight cultures started with single colonies from agar plates in LB media, followed with 1:100 dilution into 2L cultures for large-scale protein purification. After cultures reached an optical density of 0.8 at 600 nm, IPTG (final concentration 0.7 mM) was added to induce protein expression. Cultures were incubated overnight at 16 °C, and cell pastes were collected after centrifugation at 4000 rpm. The cells were resuspended in a lysis buffer (pH 7.0, 1M KCl), and sonicated. Following another centrifugation, soluble protein solution was collected and kept at 4 °C. Protein was further purified using Ni-NTA affinity resin (GE Healthcare). The column was first washed with a binding buffer containing pH 7.0, 300 mM NaCl, 20 mM TRIS, and bound protein was eluted in elution buffer containing 500 mM imidazole, 300 mM NaCl, 50 mM TRIS, pH 7.5. Thrombin was then added to cleave N-terminal histidine tag followed by a reverse Ni-NTA chromatography to remove the cleaved hexa-histidine tag. The cleaved fractions containing the enzyme without any affinity tag were pooled, concentrated and applied to the S200 (Superdex 200) column as a final purification step. Concentration of the final protein solution before crystallization was measured using UV absorption at 280 nm (Nanodrop UV spectra) and the purity is determined by SDS-PAGE. Protein solution concentrated to 10 mg/ml – confirmed by Nanodrop UV spectra (280 nm).

### Crystallization

The purified protein solution was added to crystal screen conditions in Terasaki plates (BioExpress), in 1:1 (v/v) ratio and sitting drop geometry. For the quaternary complex, protein solution was first mixed with equimolar amounts of 3-hydroxypropionate, coenzyme-A, and ATP. Initial screens produced no crystals, in apo-enzyme; beta-mercaptoethanol (BME) was added in both contexts, to reduce possible disulfide bonds between oligomers. Following the BME addition, crystals were obtained in both contexts. The apo-enzyme crystallized exclusively in 0.2M MgCl_2_, 0.1M sodium citrate tribasic dihydrate (pH 5), 10% w/v PEG. To prepare for diffraction experiments at 100 K, apo-enzyme was soaked overnight in 2.5 µl of 33% glycerol cryoprotectant (v/v) and 77% (v/v) mother liquor. Following cryoprotection, crystals were flash-frozen in liquid nitrogen.

### Data Collection and processing

Synchrotron X-ray diffraction images were collected from a single crystal of Nmar_1308 using the Pilatus 6M detector at beamline BL12-2 of the Stanford Synchrotron Radiation Lightsource (SSRL), SLAC National Accelerator Laboratory in Menlo Park, CA, USA. The diffraction data were collected to 2.1 Å resolution with unit cell dimensions a= 75.87 Å b= 75.87 Å c= 207.96 Å α= 90 β= 90 γ= 120, the space group *P*3_2_ at a wavelength of 1.0000 Å and temperature 100 K. The diffraction data were processed with the XDS package for indexing and scaled by using XSCALE^32^.

### Structure determination and refinement

Apo-Nmar_1308 structure obtained from cryo-synchrotron at SSRL, was determined by using the automated molecular replacement program PHASER^33^ implemented in PHENIX software^34^ with the previously published structure as a search model and initial rigid body refinement (PDB ID: 2DUB). After simulated-annealing refinement, individual coordinates and TLS parameters were refined. Potential positions of altered side chains and water molecules were checked by using the program COOT^35^. Then, positions with strong difference density were retained. The Ramachandran statistics for cryo-synchrotron structure (most favored / additionally allowed / disallowed) were 94 / 5 / 0% respectively. Monomer-monomer superposition was performed with the similar structures of Nmar_1308 and root-mean-squared deviations (RMSDs) were calculated based on C-alpha atom positions. Average RMSD values were calculated by taking the average of each pairwise alignment RMSD between one chain of Nmar_1308 and another from the superposed homologous structure (Fig. 4). For the overall RMSD, the average RMSDs for each chain is averaged again. Standard errors (SEs) for the average RMSDs were calculated from the standard deviations of the pairwise RMSD values between two proteins using SE = (standard deviation of pairwise RMSDs) / sqrt (N), N is the number of monomer pairs used, 6×6 =36 for hexamers and 6×3 = 18 for trimers. SE values range from 0.0003 Å to 0.008 Å. This average RMSD value between different monomers within the Nmar_1308 hexamer range from 0.16 to 0.24 A, substantially smaller than those between Nmar_1308 and the superposed homologous structures. Sequence alignment was performed by using homologous sequences from different organisms via Jalview software^27^. During the alignment, ClustalW^36^ was used as an algorithm with default parameters of Jalview. All figures were generated by PyMOL^37^ (www.schrodinger.com/pymol).

## Acknowledgments

HD acknowledges support from National Science Foundation (NSF) Science and Technology Centers grant NSF-1231306 (Biology with X-ray Lasers, BioXFEL) and The Scientific and Technological Research Council of Turkey (TUBITAK) grant (118C270). HD would like to thank Michelle Young, Ritu Khurana, Lori Anne Love and Tracy Chou for their invaluable support and discussions. The work conducted by the US Department of Energy Joint Genome Institute, a DOE Office of Science User Facility, is supported under contract no. DE-AC02-05CH11231. SW and CAF acknowledge support from U.S. Department of Energy (DOE) Office of Science, Biological and Environmental Research; Stanford Precourt Institute; and SLAC Laboratory Directed Research and Development. Portions of this research were carried out at Stanford Synchrotron Light Source (SSRL) at the SLAC National Accelerator Laboratory. SSRL is supported by the U.S. Department of Energy (DOE), Office of Science, Office of Basic Energy Sciences (OBES) under Contract No. DE-AC02-76SF00515. The SSRL Structural Molecular Biology Program is supported by the DOE Office of Biological and Environmental Research and by the National Institutes of Health, National Institute of General Medical Sciences (NIGMS) (including P41GM103393).

## Author contributions

Y.Y., S.W., C.A.F., S.D., and H.D. designed the project. H.D. prepared the samples for cryo X-ray studies. H.D., E.D., B.Y., and E.A. processed the diffraction data refined the structure. Data were analyzed by E.D., B.Y., B.B.T., E.A., Y.Y., S.W., C.A.F. and H.D. The manuscript was prepared by E.D., B.Y., B.B.T., E.A., Y.Y., S.W., C.A.F. and H.D. with input from all the coauthors.

## Competing interests

The authors declare no competing interests.

Correspondence and requests for materials should be addressed to S.W., C.A.F., or H.D.

